# Structure-Based Pharmacophore Modeling, High-throughput Screening, and Molecular Dynamics Identify a Novel DrugBank-Derived HIV-1 Protease Inhibitor

**DOI:** 10.1101/2025.11.02.686148

**Authors:** Ali Sultan Khazaal Nazal, Amjed Majid Rashid Alrashedi, Lamia Abdultef Risan Al-Iessa, Mustafa Abdulkadhim Hussein Ali Rabeea

**Author notes:** **Corresponding Author:** Amjed Majid Rashid Alrashedi.

## Abstract

HIV-1 protease (PR) is a critical enzyme for viral maturation and a major antiretroviral target. Here, a structure-based pharmacophore modeling, drug repurposing, docking, and molecular dynamics (MD) was applied to discover new PR inhibitors.

A pharmacophore model was generated from the HIV-1 PR–3TL complex (PDB 3KFP) and used to screen the DrugBank library. The identified hit and reference 3TL were docked into PR, and each complex was simulated for 100 ns. Key metrics, including binding energy, RMSD, RMSF, SASA, hydrogen bonds, salt bridges, PCA were compared.

Docking predicted BAN’s binding energy (–7.51 kcal/mol) slightly better than 3TL (–7.05 kcal/mol). MD showed BAN established a dense H-bond network, including Asp25, and a highly favorable total interaction energy (–69 kJ/mol). However, BAN binding significantly increased protease flexibility. BAN-bound PR had higher backbone RMSD (0.34 nm) and RMSF (0.215 nm) than 3TL-bound (0.22 nm, 0.121 nm), and disrupted salt bridges that remained stable with 3TL. PCA revealed BAN-bound PR sampled a larger conformational space. SASA changes were minor in all systems.

BAN binds HIV-1 PR with affinity comparable to 3TL but via a distinct mechanism, stronger polar interactions accompanied by greater protein flexibility. These results, supported by recent literature, suggest BAN as a novel scaffold for PR inhibition. Experimental validation of BAN’s inhibitory activity is warranted.

## Introduction

Human immunodeficiency virus type 1 (HIV-1) protease (PR) is a homodimeric aspartyl protease whose catalytic dyad (Asp25/Asp25′) and mobile flaps mediate cleavage of viral polyproteins into mature proteins [1,2]. Blocking this cleavage with protease inhibitors (PIs) halts virion maturation and dramatically improved AIDS outcomes [1]. Indeed, ten PIs like darunavir and lopinavir are FDA-approved for HIV therapy [2,3]. However, rapid viral mutation leads to multi-drug resistant (MDR) HIV-1 strains, limiting long-term PI efficacy [4–7]. Thus there is an urgent need for novel HIV-1 PR inhibitors, ideally identified by rapid, cost-effective methods.

Structure-based drug design (SBDD) has a proven history in HIV-1 PR inhibitor development [8]. Early peptidomimetic PIs were derived from crystal structures of PR with substrates and inhibitors, demonstrating the power of structural data in lead optimization [1,8]. Modern computational techniques, including pharmacophore modeling [9,10], [11,12], [13,14], and molecular dynamics (MD) [11,15] have increasingly enabled the exploration of vast compound libraries and in silico evaluation of binding at atomic detail [16,17]. High-throughput docking of compound libraries, including FDA-approved drugs or fragment libraries against HIV-1 PR has successfully identified new scaffolds. In this regard, a recent study screened thousands of PubChem compounds via pharmacophore filtering and rigid-receptor docking to uncover novel hits with comparable or better predicted affinity than approved Pis [16,18].

In these computational pipelines, protein–ligand docking provides an initial binding score and pose prediction, while MD simulations validate and refine these complexes by sampling conformational flexibility and estimating binding energetics over time [16,18]. Key metrics from MD, such as root-mean-square deviation (RMSD), root-mean-square fluctuation (RMSF), solvent-accessible surface area (SASA), hydrogen-bond occupancy, salt-bridge persistence, and principal component (PCA) analyses, quantify binding stability, protein mobility, and dominant motions. For HIV-1 PR, MD studies have revealed that stable inhibitors typically lock the flaps in closed conformations and maintain strong polar contacts with catalytic residues [16,18]. In contrast, weak or atypical binders can allow greater flap mobility and reorganize intramolecular interactions. Recent efforts have even applied AI and machine learning to prioritize candidate molecules before docking, underscoring the synergy of modern computational methods [17,19].

Here we combined structure-based pharmacophore modeling, DrugBank virtual screening, molecular docking, and extended MD simulations to discover and evaluate a DrugBank-derived HIV-1 PR ligand relative to a known reference inhibitor, 3TL. The goal was to interpret the binding energetics and dynamics in the context of current literature. We analyze RMSD/RMSF convergence, SASA, ligand–protein interaction energy, hydrogen bonding patterns, intramolecular salt bridges, and principal components to assess binding stability and conformational effects. By comparing our findings to recent studies, we aimed to explain how the novel hit engages HIV-1 PR and how it differs from the reference inhibitor, thereby guiding future optimization.

## Materials and methods

### Data gathering of preprocessing

HIV-1 Protease (PR), complexed with a reference inhibitor, 3TL or as we call it from now on ‘*3TL*’ was obtained from the Protein Data Bank (PDB; https://www.rcsb.org/) using the PDB ID 3KFP. This structure was derived from a fragment-based crystallographic screen of wild-type NL4-3 HIV Protease [20]. The crystal structure was prepared for computational analysis as described before [21–26]. Accordingly, all non-essential molecules, including water molecules, co-solvents, and ions, were removed from the PDB file. The coordinates of the PR monomer and the co-crystallized inhibitor/fragment were retained. Hydrogen atoms and appropriate bond orders were added to the polar residues of protein structure and the bound ligand using UCSF Chimera and Avogadro, respectively. The prepared protein structure was used to define the binding site for subsequent pharmacophore modeling, molecular docking and dynamic simulation.

### Structure-based pharmacophore nodel generation

A structure-based pharmacophore model was generated by analyzing the specific non-covalent interactions between the key reference inhibitor3TL and the HIV-1 Protease residues. Accordingly, LigandScout software v4.4.7 software was used for preparing the pharmacophore as described before [9]. Briefly, the 3D coordinates of the prepared HIV-1 PR–inhibitor complex (PDB ID 3KFP) were imported into LigandScout. The software was employed to automatically identify and annotate essential pharmacophoric features, which typically include Hydrogen Bond Acceptors (HBA), Hydrogen Bond Donors (HBD), Hydrophobic features, and Aromatic Ring features, based on the physicochemical properties and spatial arrangement of the co-crystallized ligand and the surrounding amino acid residues in the binding pocket. The automatically generated pharmacophore model was manually inspected and refined to ensure chemical relevance and maximize selectivity. Accordingly, features contributing to critical anchor points, such as those forming hydrogen bonds with the backbone or conserved catalytic residues like Asp25/Asp25’ in HIV PR), were retained.

### Preparation of chemical library and screening

The selected pharmacophore model was utilized to perform a high-throughput virtual screening. In this regard, a comprehensive compound library of DrugBank database was prepared for screening based on the previously reported study [10]. Compounds were subjected to conformational sampling and energy minimization using LigandScout’s integrated features with the Omega best algorithm to generate a set of energetically favorable 3D conformers for each molecule [9].

The generated pharmacophore model was used as a spatial filter to screen the prepared DrugBank library. The process identified compounds whose conformers could be successfully mapped onto all of the pharmacophoric features in the model. The screened compounds were ranked based on their Pharmacophore-Fit Score, which quantifies how well the compound’s features map spatially onto the pharmacophore query. The top-scoring compounds, along with the reference compounds, were designated as “*hit*” compounds for further analysis, such as molecular docking and molecular dynamics simulations.

### Molecular docking

Molecular docking studies were conducted to validate the predicted binding affinity and conformation of the reference and pharmacophore-matched hit compounds within the HIV-1 PR active site. Docking was performed using AutoDock Vina v1.2.3 [27,28] via the UseGalaxy server (accessible at https://usegalaxy.eu) [29,30]. The receptor (HIV-1 PR) was treated as rigid, and the hit ligands, in their energy-minimized 3D conformations, were treated as flexible. A grid box parameter was defined using MGLTools v1.5.7 to encompass the entire pharmacophore features obtained as described abode visually. The Vina scoring function was used to generate nine distinct docking poses for each reference 3TL and hit(s) compounds. The final lead compounds were prioritized for molecular dynamic simulation.

### Molecular dynamics simulations

Molecular dynamics (MD) simulations of HIV protease apo, and holo systems, including protease complexed with 3TL and the hit compoud were carried out using the GROMACS v2022 software package with the AMBER99SB force field [31]. The Ligands’ topology was generated using the AnteChamber Python Parser (acpype) package Version 2021.02.05.22.15 [32]. The parameterization utilized the General Amber Force Field Version 21.10 (GAFF) for describing intra-molecular interactions. Atomic partial charges were derived using the AM1-BCC (Austin Model 1-Bond Charge Correction) method. This two-step approach first calculates the electrostatic potential-derived charges using the semi-empirical AM1 Hamiltonian, followed by empirical bond charge corrections to ensure charges accurately reflect molecular polarity [33–35]. The initial protein structure was embedded in a cubic periodic box with a minimum distance of 1.0 nm between the solute and the box boundary. The system was solvated with 10,139 water molecules using the TIP3P water model. To ensure charge neutrality, an appropriate number of counter-ions (Na^+^/Cl^−^) were added.

Prior to dynamics, energy minimization was performed using the steepest descent algorithm until the system reached a maximum force below 1000 kJ mol^−1^, removing steric clashes and unfavorable contacts. Following energy minimization, the systems were equilibrated in two successive phases. First, a constant-volume (NVT) ensemble was applied for 50 ps using a leap-frog integrator with a 2 fs timestep, during which the system temperature was maintained at 300 K with a V-rescale thermostat applied separately to protein and solvent groups (τ = 0.1 ps). All bonds were constrained using the LINCS algorithm, and electrostatic interactions were treated with the Fast smooth Particle Mesh Ewald (PME) method with a 1.0 nm real-space cutoff, while van der Waals interactions were truncated at 1.0 nm using the Verlet scheme. This was followed by 50 ps of constant-pressure (NPT) equilibration, employing the Parrinello–Rahman barostat with isotropic coupling at 1 bar (τp = 2.0 ps, compressibility = 4.5×10^−5^ bar^−1^). During both phases, periodic boundary conditions were applied in all three dimensions, and long-range dispersion corrections were included for energy and pressure. Following equilibration, a 100 ns production MD simulation was performed under periodic boundary conditions in all three dimensions. Electrostatics were computed using the Fast smooth Particle Mesh Ewald (PME) electrostatics method with a real-space cutoff of 1.0 nm, while van der Waals interactions were treated using the Verlet cutoff scheme, with long-range dispersion corrections applied to both energy and pressure. Coordinates were saved every 10 ps for subsequent analyses.

### Trajectory analysis

To statistically evaluate the conformational stability and dynamics of the simulated systems [21], key metrics were calculated from the convergence production phase of the trajectories. For all time-series data the analysis included the Mean and Standard Deviation (StDev) to assess central tendency and spread, the Minimum (Min) and Maximum (Max) values to describe the range of motion, and the Standard Error of the Mean (SEM) to quantify the precision of the mean estimate. Furthermore, to focus specifically on the stable molecular dynamics, the Production Mean (ProdMean) and Production Standard Deviation (ProdStDev) were calculated using the last 80% of the simulation time. For the residue-specific data (RMSF), which is not a time-series, the overall Mean, StDev, SEM, Min, and Max were calculated across all residues to quantify the average and range of local flexibility. All statistical results were compiled and presented in a structured text file for direct comparison.

### Hydrogen bond analysis

Hydrogen bond (H-bond) analysis was performed using the Hydrogen Bonds plugin of Visual Molecular Dynamics (VMD). To ensure accurate detection of hydrogen bonds throughout the simulation, the following parameters, including all frames of the 100 ns trajectory, donor-acceptor distance cutoff equal to 3.0 Å, angle cutoff of 20°, selection type of both donors and acceptors, enabaled update selections every frame, and disabled polar atoms restriction (all atoms considered) were considered.

### Principal component analysis

PCA was performed to characterize the essential motions of the system throughout the molecular dynamics (MD) simulation. The covariance matrix of atomic positional fluctuations was calculated after least-squares fitting of the trajectory to the reference structure to remove overall translational and rotational motions. Eigenvalues and eigenvectors were obtained by diagonalizing the covariance matrix using GROMACS tools, and the eigenvalues were used to estimate the contribution (variance explained) of each principal component (PC) to the total motion. The first four PCs, were used for further analysis. The percentage of variance explained by each PC was calculated as [36]:

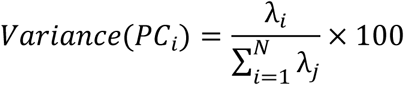

where λi is the eigenvalue of the iii-th PC and N is the total number of components.

#### Salt bridge analysis

Salt bridge interactions were analyzed to evaluate the stability and persistence of electrostatic contacts during the molecular dynamics (MD) simulation. The analysis was performed using the Salt Bridges Plugin (version 1.0) implemented in VMD over the entire trajectory. Protein residues were selected as the target atom group, and salt bridges were identified based on an oxygen–nitrogen distance cutoff of 3.2 Å, with selections updated for each frame. The descriptive statistics, including the mean, minimum, maximum, and standard deviation of distances, as well as the occupancy percentage, defined as the fraction of trajectory frames in which the oxygen–nitrogen distance was ≤ 3.2 Å.

## Results

### Pharmacophore-based screening

The pharmacophore extraction of 3TL in its complex with the HIV-1 protease resulted in eight pharmacophore features including, two hydrophobic regions, two H-bond acceptors, four H-bond donors sites. Virtual screening of the Drugbank library with 9.2K molecules has resulted in identification of one potent hit with a generic name of HONH-BENZYLMALONYL-L-ALANYLGLYCINE-P-NITROANILIDE (DrugBank Accession Number: DB07434). The relative pharmacophore-fit score was 0.93, matching all 3TL-pharmacophore extracted features (Figure 1). The compound, which from now on we call it ‘*BAN’* through the manuscript, was composed of 200 conformations that it 40^th^ was match with all pharmacophore features with the energy of 98.344.

**Fig 1.**
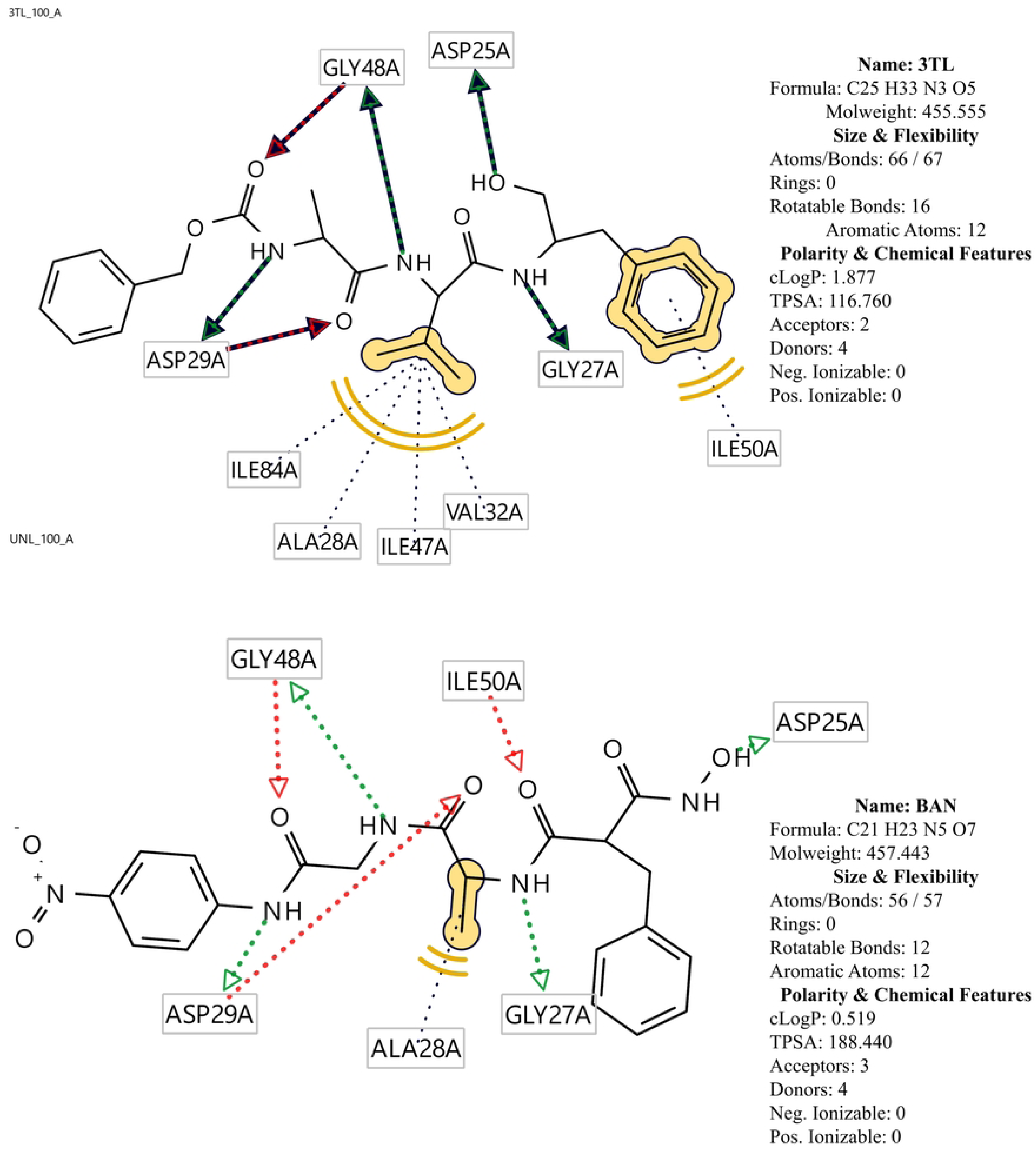
Matching of two reference 3TL compound and the hit Ban compound within the pharmacophore in the HIV-1 protease. Near aminoc acid residues are depicted. Yellow spears are hydrophobic regions, Green and Red arrows are H bond donors and acceptors, respectively

### Computational affinity measurements

Molecular docking of the identified hit compound BAN and the reference HIV-1 protease inhibitor 3TL was performed using AutoDock Vina to evaluate their binding affinities and interaction profiles within the protease active site. The docking results are summarized in terms of binding energy (Vina score), intermolecular and intramolecular contributions, unbound energy, and torsional flexibility. The binding affinity predicted for BAN was –7.51 kcal·mol⁻¹, which is slightly more favorable than the reference inhibitor 3TL (–7.05 kcal·mol⁻¹). This result indicates that BAN is capable of achieving a binding energy comparable to the established inhibitor, suggesting potential for inhibitory activity against HIV-1 protease. The difference, while modest (∼0.46 kcal·mol⁻¹), may reflect variations in the electrostatic stabilization and conformational flexibility of the two ligands.

The decomposition of binding energies revealed that BAN exhibited an intermolecular interaction energy (INTER) of –9.75 kcal·mol⁻¹, comparable to 3TL (–9.69 kcal·mol⁻¹). This suggests that both ligands establish similar levels of stabilizing contacts with the protease binding pocket. However, BAN demonstrated a slightly reduced intramolecular strain (INTRA: –3.41 kcal·mol⁻¹) compared to 3TL (–3.84 kcal·mol⁻¹), implying that BAN adopts its bound conformation with less energetic penalty. The lower strain energy may contribute to its marginally stronger overall binding score. Unbound energy values were also lower for BAN (–1.04 kcal·mol⁻¹) compared with 3TL (– 1.35 kcal·mol⁻¹), reinforcing the observation that BAN achieves stable binding with minimal conformational destabilization relative to its free state.

Analysis of torsional flexibility showed that BAN possessed 12 active torsions, whereas 3TL exhibited 16 active torsions. The reduced number of rotatable bonds in BAN indicates a lower degree of conformational entropy loss upon binding, which may favor its docking score despite its smaller molecular complexity compared to 3TL. Importantly, this suggests that BAN is able to maintain binding efficacy while imposing a lower entropic cost on the system. These findings highlights the need for a deeper investigation into highlightin BAN as a potent inhibitor of HIV-1 protease through molecular dynamic simulation as of follow.

### MD simulation

The stability and dynamics of the target protein, both in its apo form Apo-protease and in complex with ligands 3TL and BAN, Holo systems, were evaluated over a 100 ns simulation period. Analyzes focused on HIV-1 protease stability with RMSD, local residue flexibility with RMSF, protein surface area changes (SASA), specific ligand-protein affinity or interaction energy, hydrogen binding, salt-bridge and PCA analysis.

#### Conformational stability and overall trajectory convergence

The root mean square deviation (RMSD) of the protein backbone was monitored to assess the structural stability and convergence of the three simulated systems (Figure 2). The Holo 3TL system exhibited the highest stability and fastest convergence. After an initial slight adjustment in the first 20 ps, the system rapidly stabilized, maintaining a low average backbone RMSD of ≈0.223 nm during the production phase (>20 ps). The low standard deviation of σ=0.026 nm (Table 1) indicates minimal structural fluctuation relative to the starting conformation.

**Fig 2.**
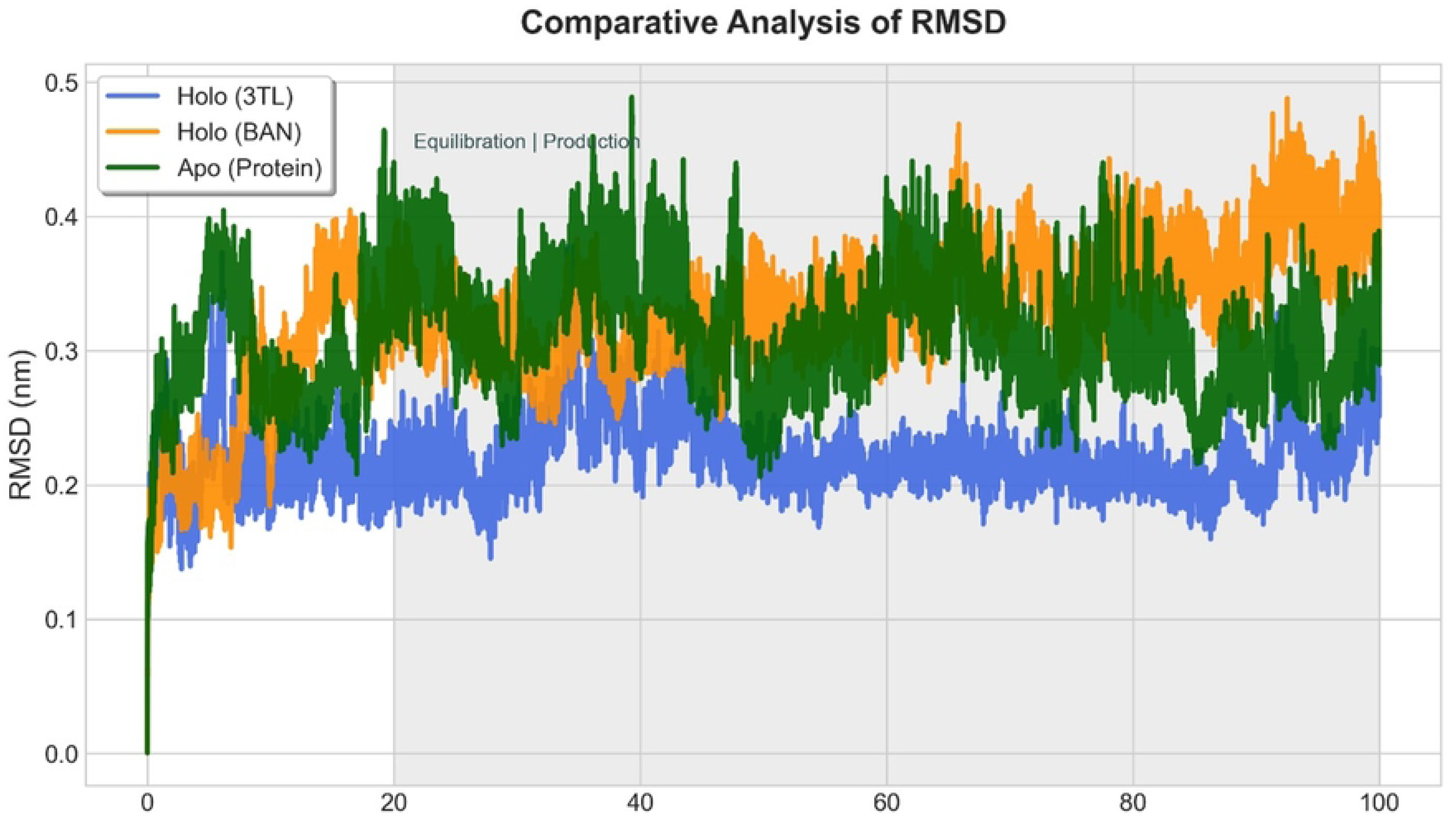
Comparative analysis of protein backbone RMSD. RMSD trajectories of the protein backbone for the Apo system (Apo-Protein, dark green) and the two holo complexes (Holo (3TL), royal blue; Holo (BAN), orange) over 100 ns. The shaded region denotes the production phase (after 20 ns), used for calculating convergence statistics. The Holo (3TL) system demonstrated the lowest average deviation (μ=0.223 nm), indicating the highest conformational stability

**Table 1.** Descriptive statistics of RMSD findings.

| **Metric (nm)** | **Holo (3TL)** | **Holo (BAN)** | **Apo (Protease)** |
| --- | --- | --- | --- |
| Mean | 0.221 | 0.327 | 0.313 |
| ProdMean | 0.223 | 0.342 | 0.318 |
| Max | 0.378 | 0.488 | 0.489 |
| Min | 0.001 | 0.001 | 0.001 |
| ProdStDev | 0.026 | 0.037 | 0.041 |
| SEM | 0 | 0 | 0 |
| StDev | 0.027 | 0.052 | 0.044 |

The Holo BAN system demonstrated higher fluctuation and less stability compared to the 3TL complex, oscillating between 0.25 nm and 0.45 nm throughout the production phase. The mean RMSD was substantially higher at 0.342 nm (σ=0.037 nm). This larger deviation suggests that the binding of the BAN ligand induces a greater conformational change or results in a less rigid final complex structure. The Apo-protease system displayed stability characteristics intermediate between the two holo systems, converging around a mean RMSD of 0.318 nm (σ=0.041 nm). Although the Apo system’s mean is lower than Holo BAN, its larger standard deviation indicates slightly broader conformational sampling. Accordingly, the Holo 3TL complex achieves the most stable conformation, suggesting the 3TL ligand is highly effective at rigidifying the protein structure compared to both the Apo form and the Holo BAN complex.

#### Local flexibility analysis

Root Mean Square Fluctuation (RMSF) was calculated per residue to identify regions of local flexibility and how ligand binding affects these dynamics (Figure 3). The RMSF plots for the Apo-protease and Holo 3TL complex trajectories were remarkably similar, with a low overall mean RMSF of 0.137 nm and 0.121 nm, respectively (Table 2). This indicates that the 3TL ligand binds without drastically altering the intrinsic flexibility profile of the Apo protein. In contrast, the Holo BAN system exhibits significantly higher RMSF values across multiple loop regions, particularly around the active site or domain interfaces (residue indices ≈50−250 and ≈750−950, and a large peak near residue 1500). The mean RMSF for Holo BAN was nearly double that of the other two systems (0.215 nm), and its maximum fluctuation reaches 0.874 nm. The elevated flexibility in the Holo BAN complex confirms that the ligand induced protease instability as observed in the RMSD data. The BAN ligand appears to destabilize critical regions of the protein, potentially due to transient interactions or an unstable binding mode.

**Fig 3.**
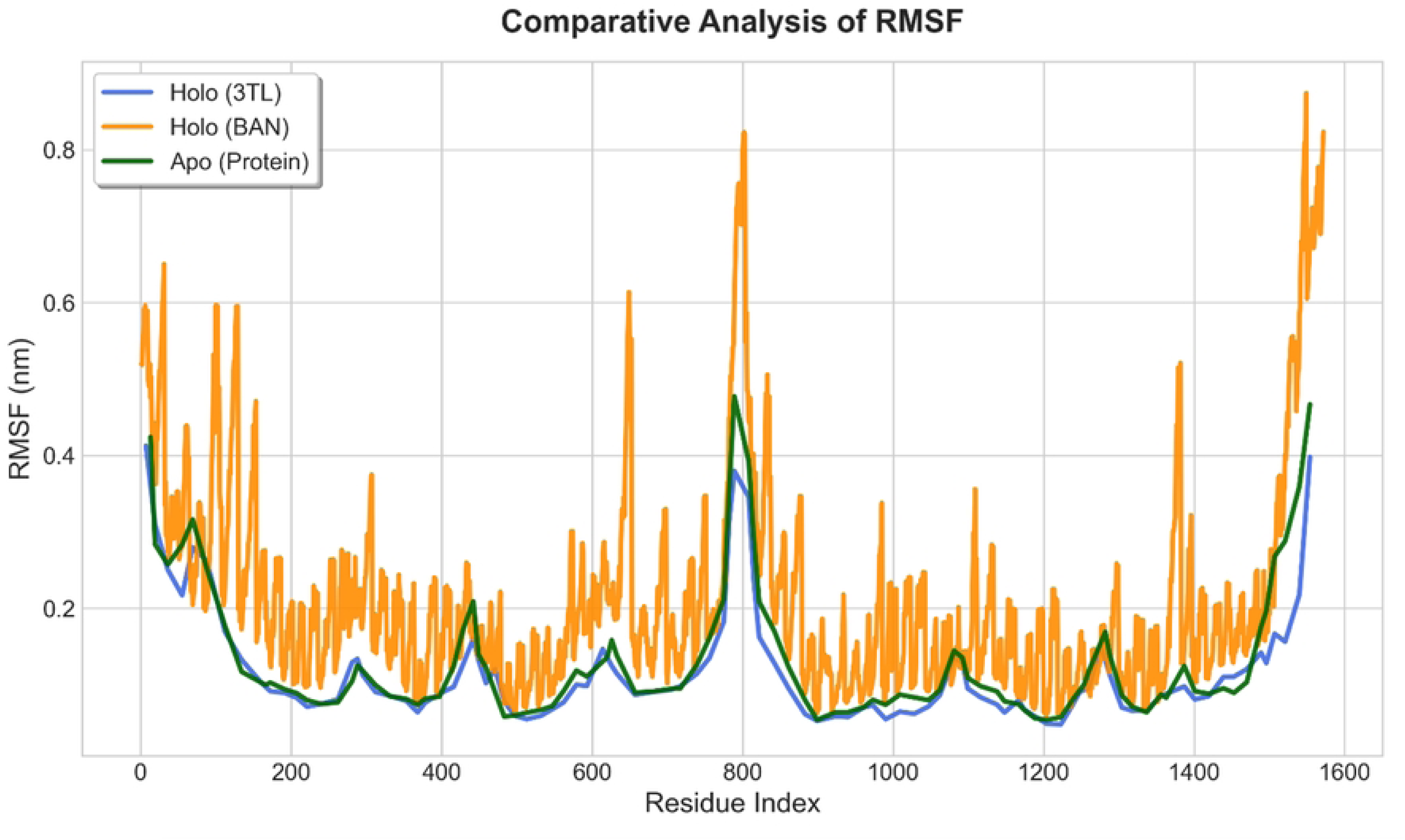
Comparative analysis of residue-wise RMSF. Root Mean Square Fluctuation (RMSF) per residue for the Apo system (dark green) and the Holo complexes (3TL in blue, BAN in orange). The Apo-Protein and Holo (3TL) systems share a similar, low-fluctuation profile. The Holo (BAN) complex exhibits significantly elevated flexibility (higher peaks) across several loop regions, confirming structural destabilization induced by the BAN ligand

**Table 2.** Descriptive statistics of RMSF findings.

| Metric (nm) | Holo (3TL) | Holo (BAN) | Apo (Protease) |
| --- | --- | --- | --- |
| Mean | 0.121 | 0.215 | 0.137 |
| Max | 0.412 | 0.874 | 0.477 |
| Min | 0.048 | 0.06 | 0.054 |
| SEM | 0.008 | 0.004 | 0.009 |
| StDev | 0.077 | 0.141 | 0.093 |

#### Ligand-protein interaction energy

The total interaction energy provides a thermodynamic measure of the favorable or unfavorable binding affinity between the ligand and the protein. Since the interaction energy is undefined for the Apo system, this analysis was restricted to the two Holo complexes (Figure 4). As a result, the Holo BAN complex maintained highly favorable, negative, total interaction energies throughout the production phase, settling at a mean value of −69.112 kJ/mol (Table 3). This negative value signifies strong, persistent, and energetically favorable non-covalent interactions (electrostatic and van der Waals) between the BAN ligand and the receptor. Conversely, the Holo 3TL complex consistently displayed highly unfavorable total interaction energies, with a production mean of +95.529 kJ/mol. This positive energy suggests strong repulsive components, likely unfavorable electrostatic or desolvation penalties, that overwhelm any favorable van der Waals contributions. Accordingly, structurally more stable Holo 3TL system with the highly positive interaction energy indicates a thermodynamically unfavorable ligand-protein association, and the protase remains resemble to the apo system. The Holo BAN complex, on the other hand, is held together by significant and favorable binding interactions, highlighting stable bonding and altering flexibility of HIV-1 protease.

**Fig 4.**
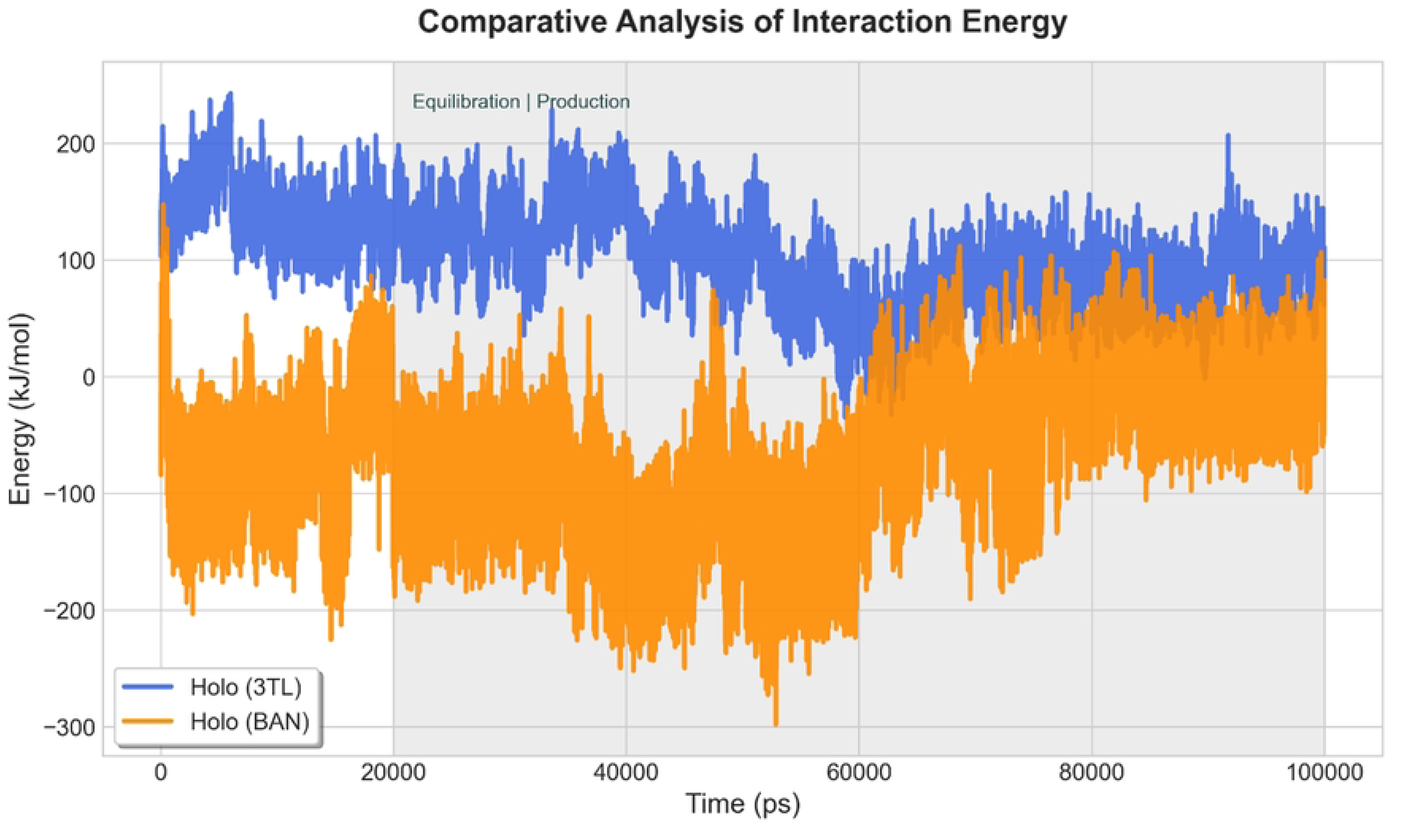
Comparative analysis of total ligand-protein interaction energy. Total non-bonded interaction energy (electrostatic plus van der Waals) between the protein and the bound ligands over 100 ns. This metric is only applicable to the holo systems. The Holo BAN complex displays highly favorable (negative) binding energy (μ=−69.112 kJ/mol), while the Holo 3TL complex shows highly unfavorable (positive) energy (μ=+95.529 kJ/mol)

**Table 3.**
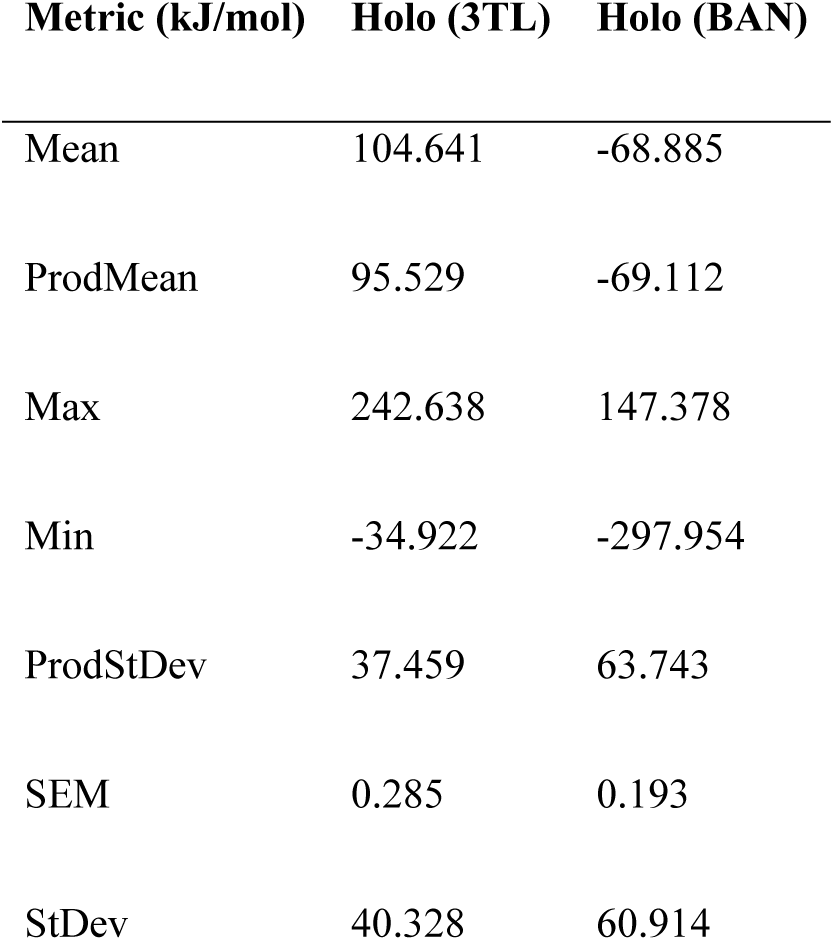
Descriptive statistics of interaction energies.

| Metric (kJ/mol) | Holo (3TL) | Holo (BAN) |
| --- | --- | --- |
| Mean | 104.641 | -68.885 |
| ProdMean | 95.529 | -69.112 |
| Max | 242.638 | 147.378 |
| Min | -34.922 | -297.954 |
| ProdStDev | 37.459 | 63.743 |
| SEM | 0.285 | 0.193 |
| StDev | 40.328 | 60.914 |

#### Solvent accessible surface area

The Solvent Accessible Surface Area (SASA) was analyzed to track conformational changes related to solvent exposure, which is often indicative of protein compaction or unfolding. All three systems exhibited similar average SASA values, with Holo 3TL at 64.314 nm^2^, Apo-Protein at 63.261 nm^2^, and Holo BAN at 65.899 nm^2^ during production (Figure 5, Table 4). The standard deviations are also tightly clustered (≈1.7−1.8 nm^2^). The data suggest that ligand binding, even by the destabilizing BAN, does not induce large-scale domain rearrangements or unfolding events that would dramatically increase or decrease the overall solvent exposure of the protein. The minor observed SASA increase in Holo BAN is consistent with the heightened residual mobility (RMSF) of external loops, which briefly expose more surface area.

**Fig 5.**
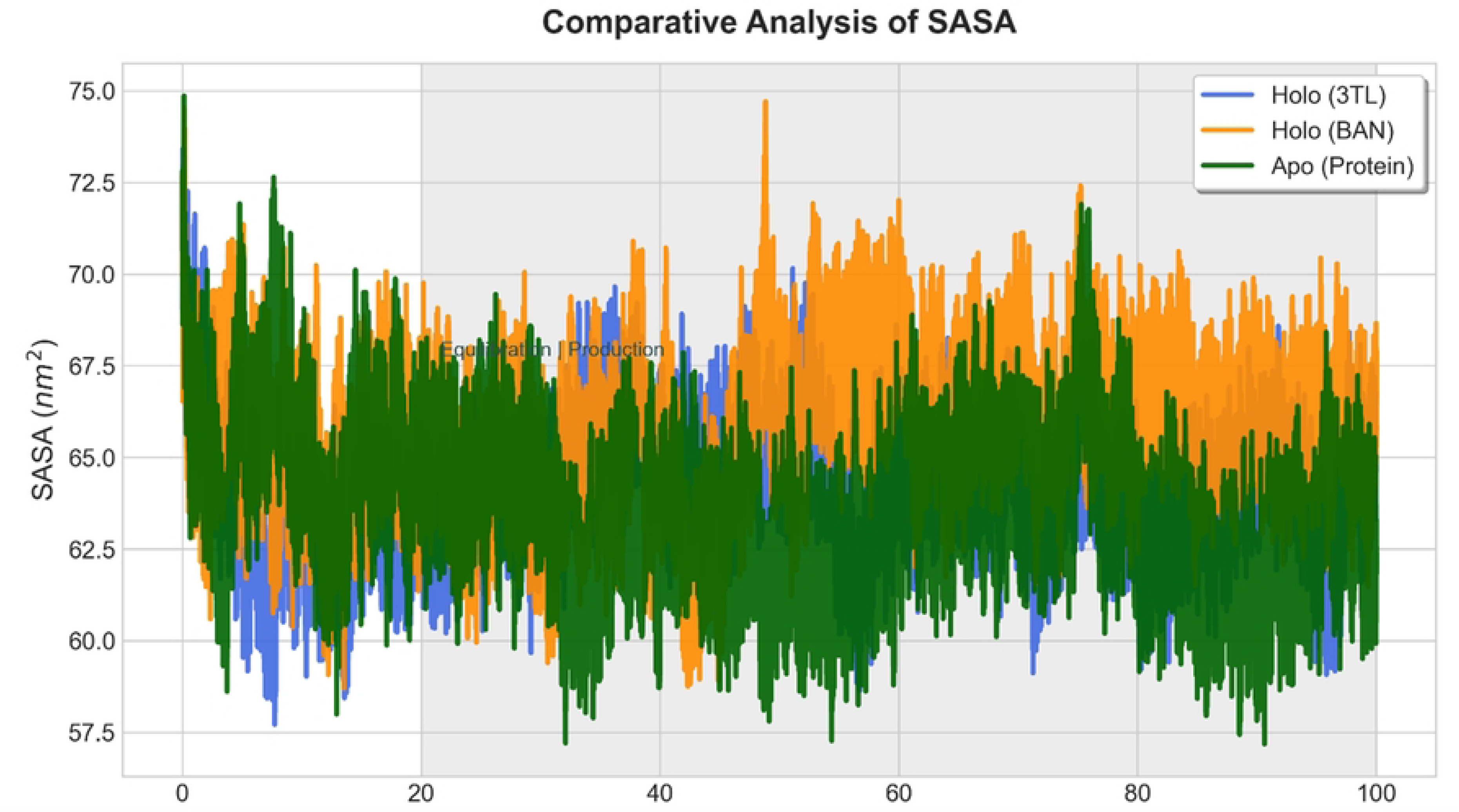
Comparative analysis of protein solvent accessible surface area (SASA). SASA trajectories for all three simulated systems over 100 ns. All systems show narrow distributions and similar mean SASA values (≈63−66 nm2), suggesting that ligand binding does not induce major changes in the overall global compaction or solvent exposure of the protein

**Table 4.** Descriptive statistics of SASA findings.

| Metric (nm<sup>2</sup>) | Holo (3TL) | Holo (BAN) | Apo (Protease) |
| --- | --- | --- | --- |
| Mean | 64.158 | 65.794 | 63.651 |
| ProdMean | 64.314 | 65.899 | 63.261 |
| Max | 73.425 | 74.703 | 74.858 |
| Min | 57.712 | 58.698 | 57.18 |
| ProdStDev | 1.637 | 1.832 | 1.761 |
| SEM | 0.013 | 0.006 | 0.006 |
| StDev | 1.799 | 1.83 | 1.992 |

#### Hydrogen bond analysis

To elucidate the binding stability and key interactions of the inhibitors with the HIV-1 protease, a detailed hydrogen bond (H-bond) analysis was performed over the course of the divergent time of MD simulation, 20 ns to 90 ns. Hydrogen bonds are critical for the affinity and specificity of ligand-protein interactions. For the reference inhibitor 3TL, a total of 13 unique hydrogen bonds were identified between the ligand and the protease active site residues. The analysis revealed several persistent interactions, although most were transient, indicating dynamic binding. The most significant H-bond was observed between the main chain of GLY48 and the side chain of 3TL, with an occupancy of 0.78%. Another notable interaction occurred between the 3TL side chain and the main chain of PHE53, showing an occupancy of 0.67%. Other residues forming transient hydrogen bonds with 3TL included MET46 (0.22%), LYS55 (0.17% and 0.11%), ILE50 (0.14%), LYS45 (0.14%), and GLY49 (0.11%). The time evolution of the total number of hydrogen bonds and the occupancy of specific interactions are detailed in Figure 6.

**Fig 6.**
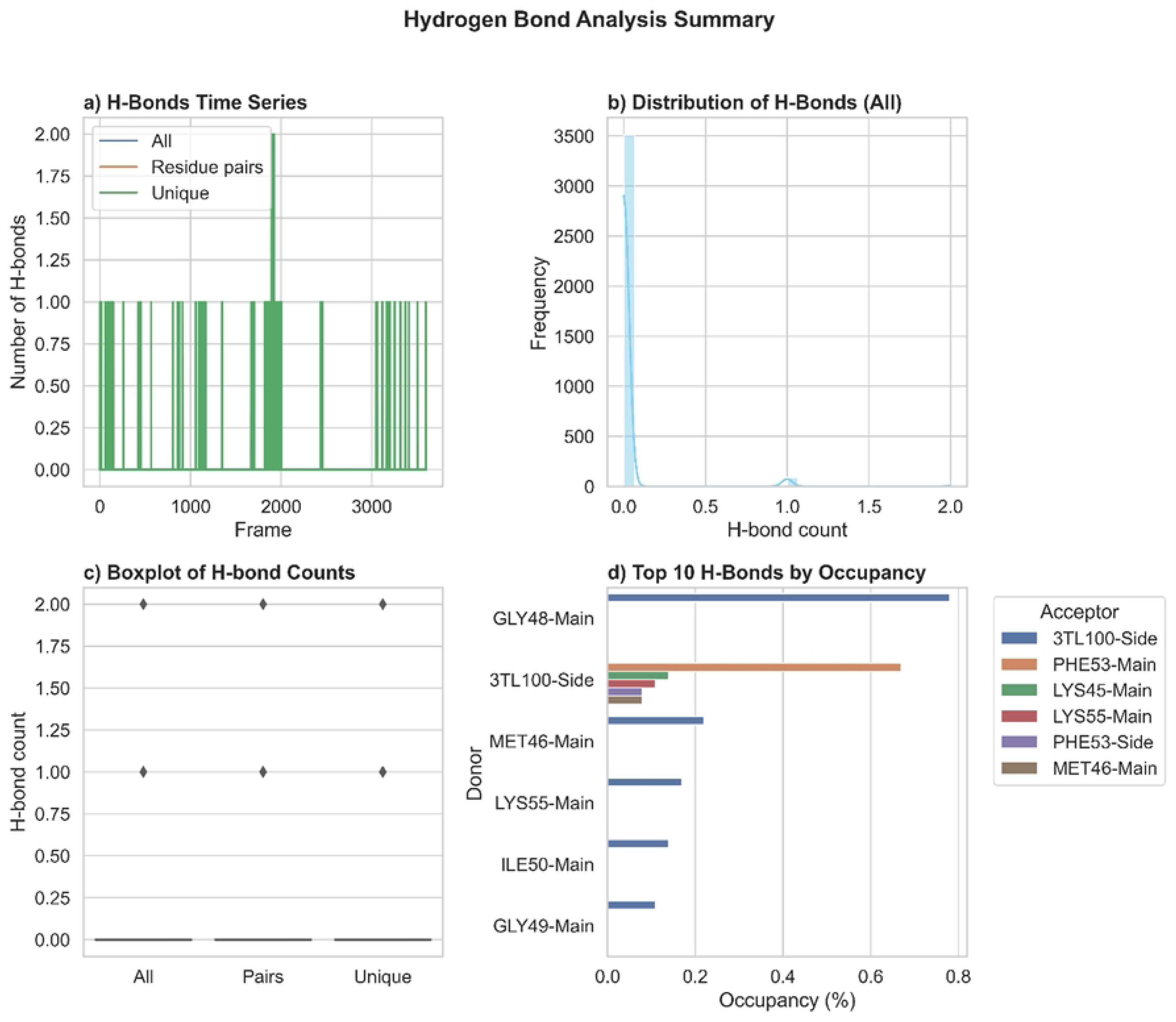
Hydrogen bond dynamics of the HIV-1 protease–3TL complex. Hydrogen bond occupancy and lifetime analysis between HIV-1 protease and 3TL over the simulation trajectory. The persistence of low hydrogen bonds emphasizes the low-to-medium polar interactions, which serve as a benchmark for comparison with the hit compound BAN

In contrast, the hit compound BAN established a more extensive and stable hydrogen bonding network, with a total of 18 unique hydrogen bonds identified (Figure 7). A remarkably persistent interaction was observed between BAN and the catalytic residue ASP25. This interaction was maintained for the majority of the simulation, demonstrating an occupancy of over 100% (165.76%), suggesting that multiple hydrogen bonds were concurrently formed between the side chains of BAN and ASP25. Furthermore, BAN formed several other stable H-bonds, including with the main chain of LEU24 (20.66%), GLY48 (11.38%), and GLY49 (8.29%). This robust network of interactions suggests a strong binding affinity of BAN for the protease, potentially greater than that of the reference inhibitor 3TL.

**Fig 7.**
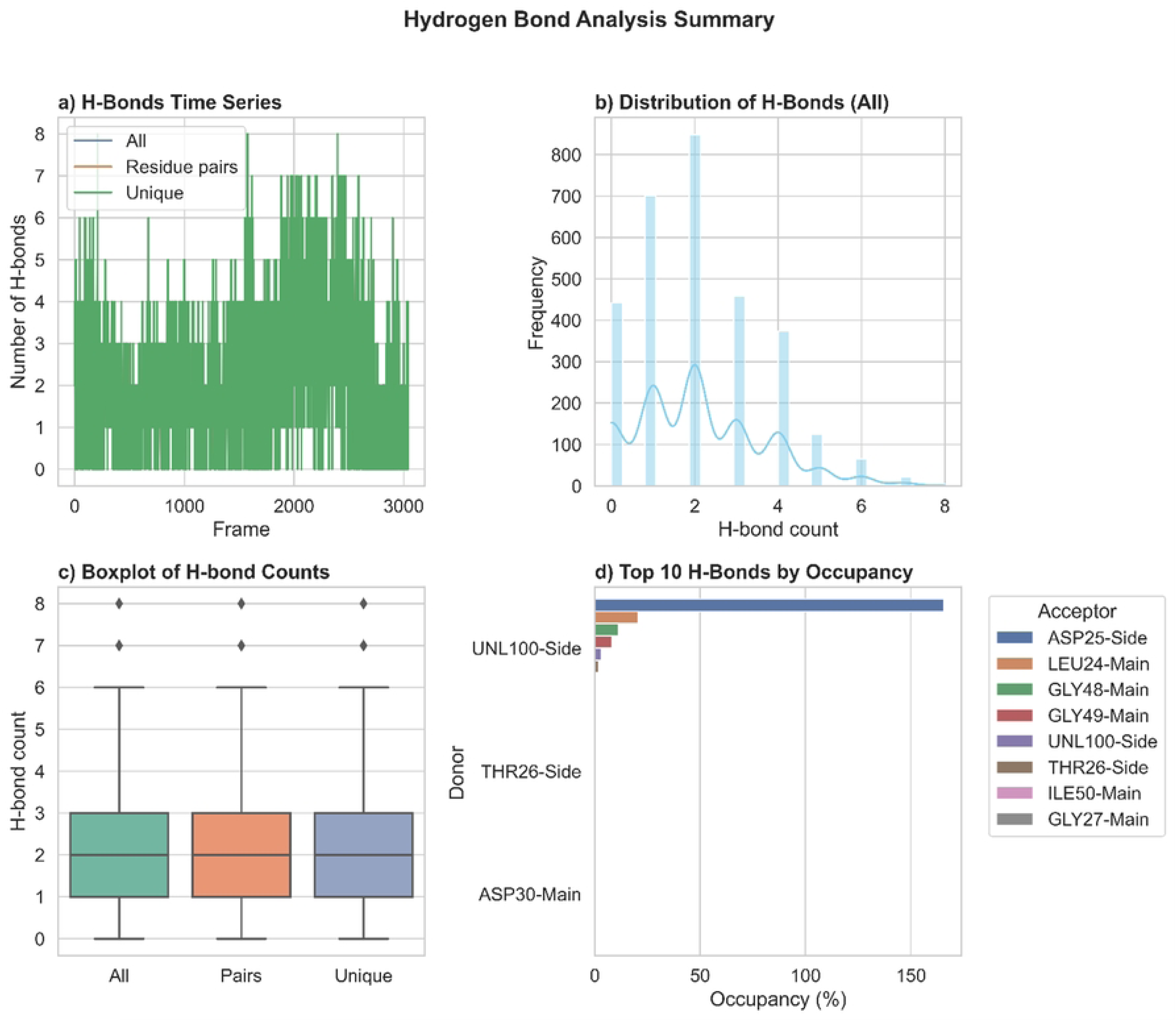
Binding conformation of BAN within the HIV-1 protease active site. Representative structural snapshot of the HIV-1 protease–BAN complex obtained from the convergenced MD simulations at 20 ns to 100 ns. The identified hit compound BAN occupies the catalytic cleft and shows close proximity to the catalytic Asp25/Asp25′ dyad. The orientation suggests potential for hydrogen bonding and hydrophobic contacts, indicating its suitability as a protease inhibitor scaffold. All paird hydrogen bindings were unique. * UNL or Unknown ligand = BAN

#### Salt bridge analysis

Salt bridges contribute significantly to the structural stability and integrity of proteins. We analyzed the intramolecular salt bridges within the HIV-1 protease structure when complexed with each inhibitor to assess conformational stability. Several key salt bridges were monitored throughout the simulation. For the 3TL complex, the most stable salt bridge was formed between GLU34 and LYS20, demonstrating an occupancy of 18.66%. This was followed by an interaction between ASP30 and LYS45, which had an occupancy of 10.24%. The majority of other potential salt bridges were found to be highly transient, with occupancies below 2%. In this regard, the interaction between ASP29 and ARG87 was present for only 0.99% of the simulation time. The persistence of these specific electrostatic interactions suggests their importance in maintaining the folded architecture of the protease during the simulation (Figure 8).

**Fig 8.**
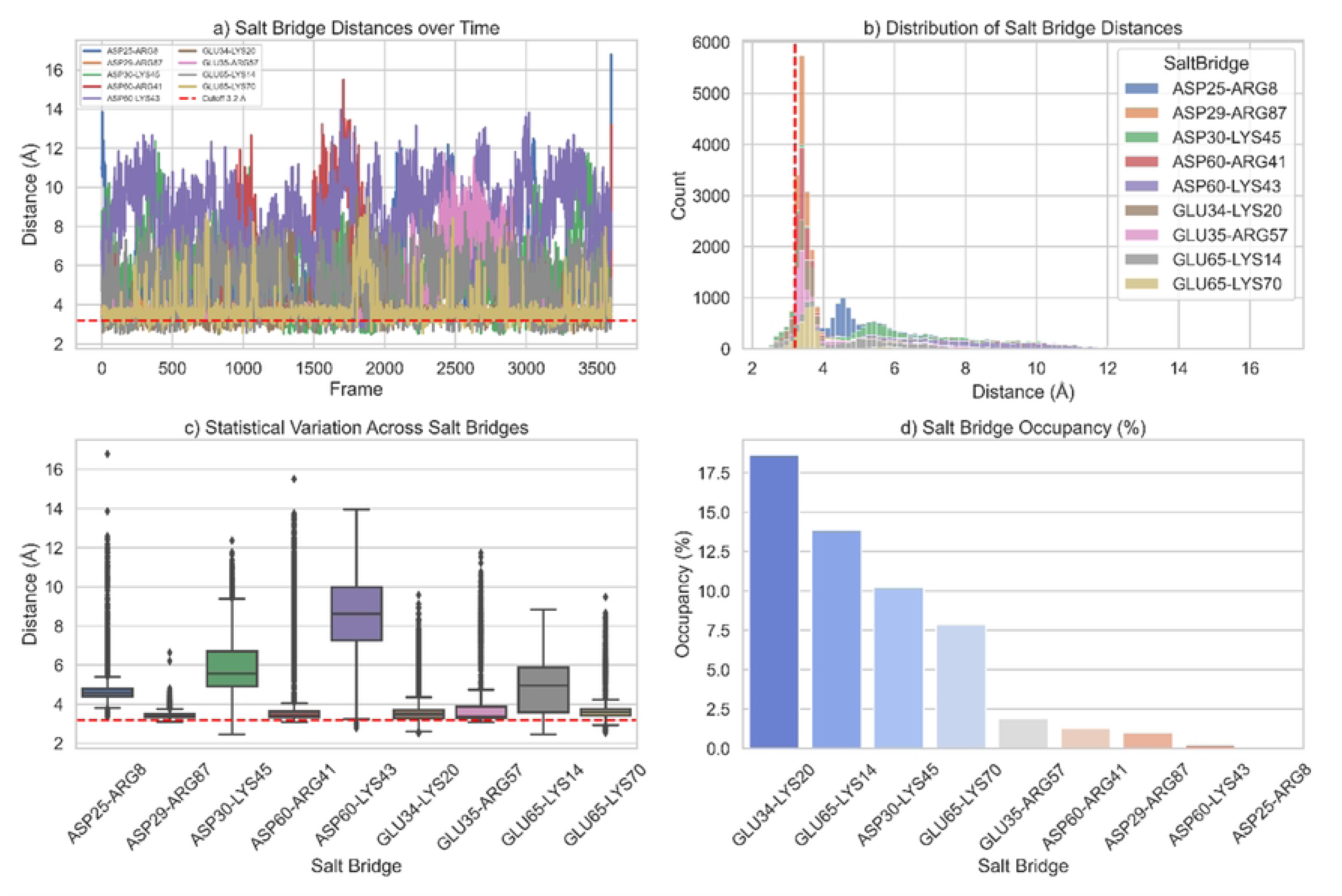
Salt bridge dynamics of the HIV-1 protease–3TL complex. Time-dependent profile of salt bridge interactions formed between charged residues of HIV-1 protease and the reference inhibitor 3TL during a 100-ns MD simulation. The plot demonstrates stable and persistent salt bridge contacts that contribute to structural stabilization of the protease–inhibitor complex, highlighting the role of electrostatic interactions in maintaining high-affinity binding

As shown in Figure 9, the analysis of the BAN-protease complex revealed a notable decrease in the stability of these key intramolecular salt bridges. The occupancy of the GLU34-LYS20 interaction dropped significantly to 3.91%, and the ASP30-LYS45 interaction was present for only 2.47% of the simulation time. This reduction in salt bridge stability suggests that the binding of BAN may induce a conformational state in the protease that is distinct from the 3TL-bound form, potentially involving greater flexibility or a slight rearrangement of charged residues.

**Fig 9.**
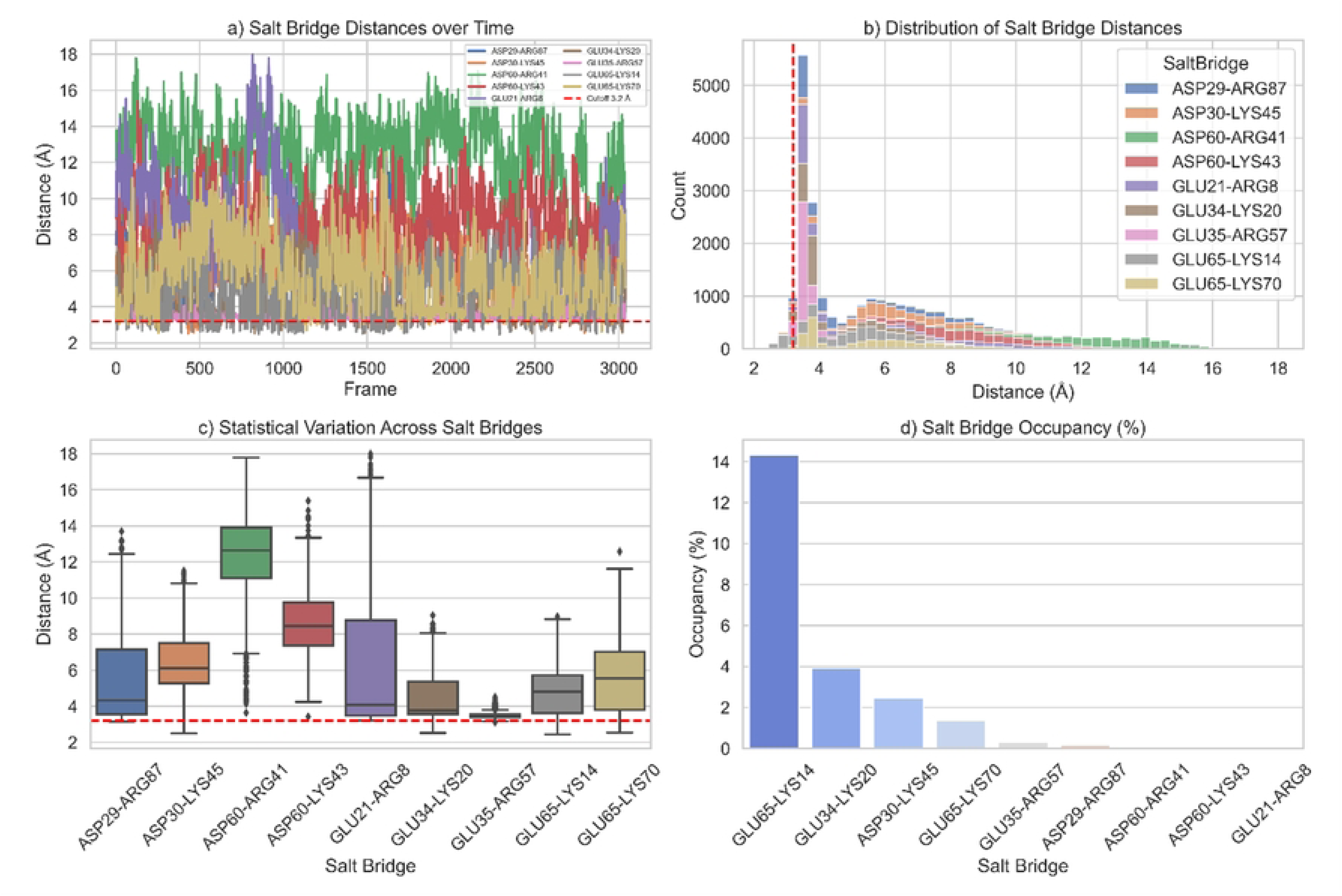
Salt bridge dynamics of the HIV-1 protease–BAN complex. Time evolution of salt bridge interactions in the BAN-bound protease. The occupancy and stability of electrostatic contacts suggest differences in charge stabilization compared to the 3TL reference complex. The data indicate that BAN binding induces a distinct electrostatic interaction pattern, potentially contributing to an altered dynamic profile of the protease

#### Principal component analysis

PCA was performed on the C-alpha atoms of the protease to identify the large-scale collective motions and explore the conformational landscape sampled by the enzyme when bound to the inhibitors. The analysis condenses the complex atomic fluctuations into a set of principal components (PCs), with the first few PCs describing the most significant protein motions. For the 3TL-protease complex, the first four principal components (PC1-PC4) accounted for 61.29% of the total variance in atomic motion observed during the simulation. PC1, representing the most dominant motion, contributed 37.02%, while PC2 contributed 10.17% of the total variance. The projection of the simulation trajectory onto the plane defined by PC1 and PC2 reveals that the protein atoms occupied a relatively defined and compact conformational space, suggesting that the binding of 3TL confers significant stability to the enzyme, restricting large-scale movements (Figure 10). The porcupine plots illustrate that the dominant motion captured by PC1 corresponds to the characteristic “flapping” motion of the protease flaps.

**Fig 10.**
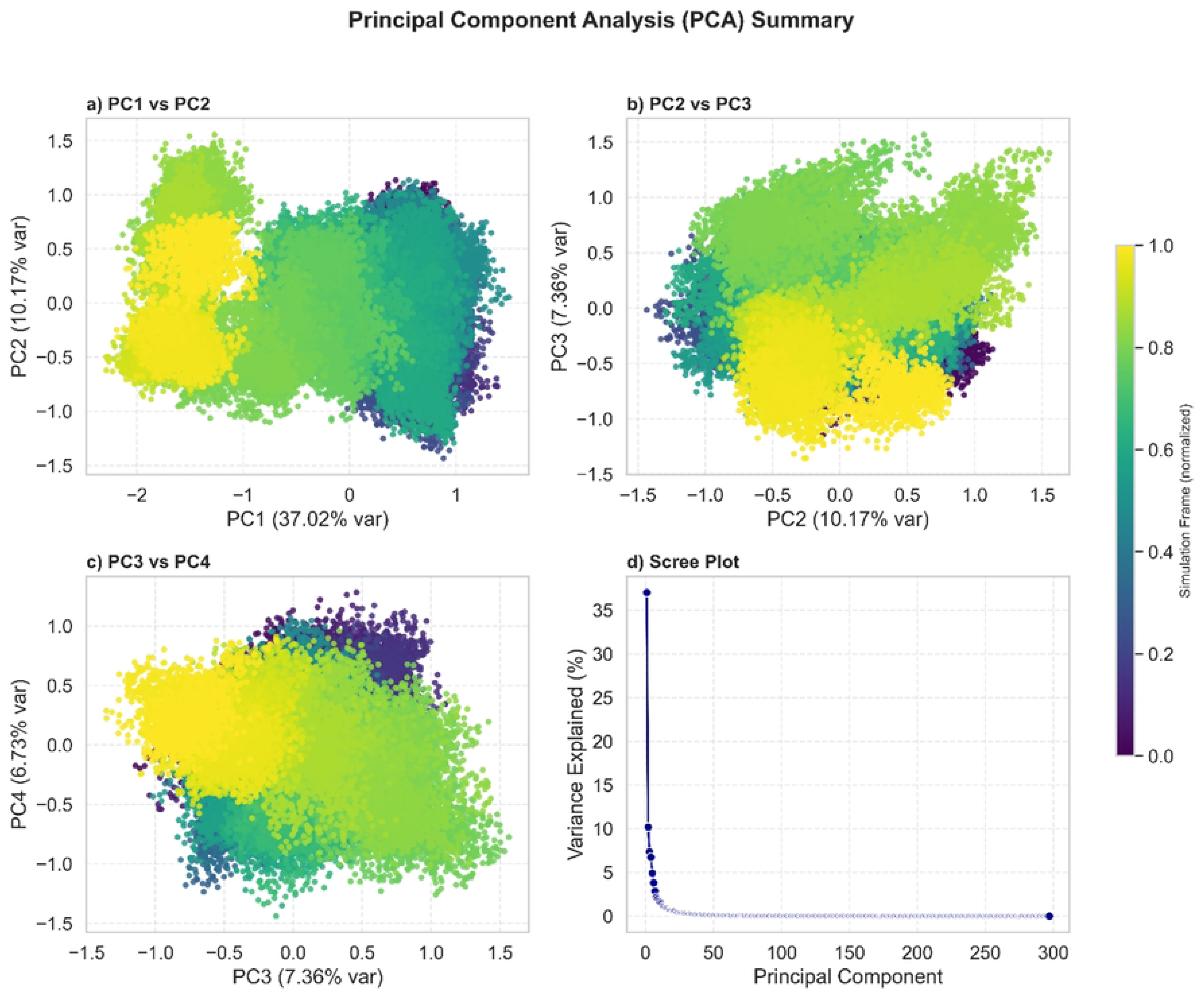
Principal component analysis of the HIV-1 protease–3TL complex. PCA plots describing the conformational landscape of the protease in complex with 3TL. (A, B, and C) Projection of trajectory frames along PC1 vs. PC2, PC2 vs. PC3, and PC3 vs. PC4 showing limited conformational fluctuations. (D) Eigenvalue distribution reflecting constrained collective motions. Free energy landscapes indicating well-defined minima corresponding to stable states of the protease in the presence of 3TL. These findings suggest that 3TL binding restricts protease flexibility more effectively than BAN

The PCA of the BAN-protease complex indicated a greater degree of conformational flexibility (figure 11). The first four PCs accounted for a larger portion of the total variance (72.07%), with PC1 alone responsible for 45.70% of the motion. The increased variance captured by the initial principal components, along with a higher overall trace of the covariance matrix (3.49 for BAN vs. 2.15 for 3TL), suggests that the protease experiences more pronounced and coordinated motions when bound to BAN. The 2D projection plot for the BAN complex is expected to cover a larger conformational space compared to 3TL, consistent with the reduced salt bridge stability and indicative of enhanced flap dynamics. This heightened flexibility could have implications for the inhibitor’s mechanism of action and residence time in the active site.

**Fig 11.**
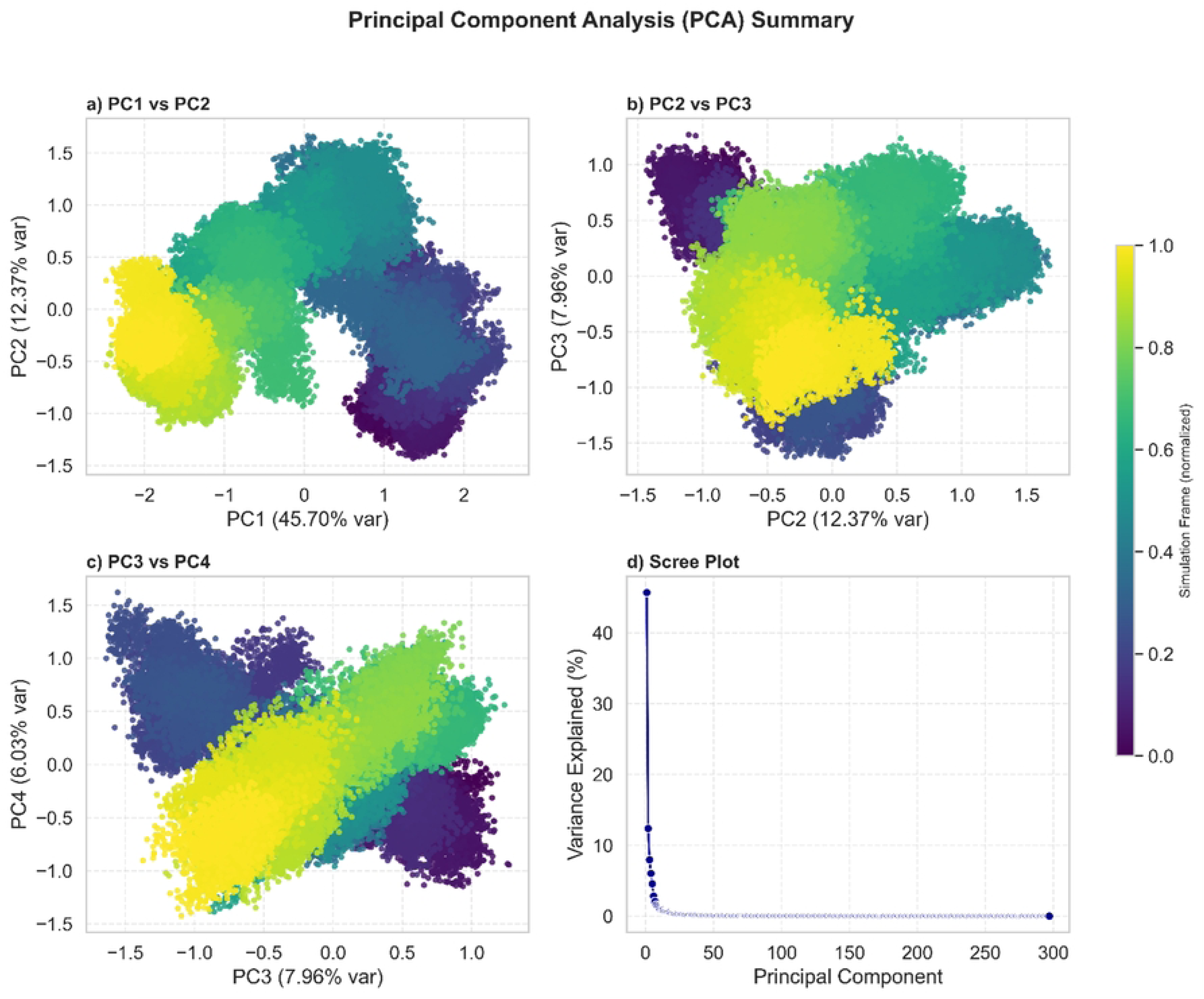
Principal component analysis (PCA) of the HIV-1 protease–BAN complex. Essential dynamics of the BAN-bound protease represented by PCA. (A, B, and C) Projection of the MD trajectory along the first two principal components (PC1 vs. PC2), showing conformational sampling. (D) Distribution of eigenvalues, indicating the dominance of low-frequency motions. Two-dimensional free energy landscapes constructed from PC1 and PC2, highlighting preferred conformational states of the protease in the presence of BAN. Together, the analysis reveals the conformational adaptability of the enzyme upon BAN binding

## Discussion

The high-throughput pharmacophore screen of DrugBank yielded BAN as a top hit matching all critical features of the 3TL pharmacophore. Docking predicted BAN’s binding energy to be slightly better than the reference 3TL. This modest advantage likely arises from BAN’s favorable intermolecular contacts and lower intramolecular strain. BAN’s interaction energy in MD was highly favorable (mean –69.1 kJ/mol), whereas 3TL’s was not. Such a positive interaction energy for 3TL suggests that any favorable van der Waals contacts were offset by desolvation or repulsive electrostatics, consistent with the ligand rigidifying the protease without forming compensating polar bonds. In contrast, BAN formed persistently favorable interactions that stabilized the complex. These observations mirror prior reports, stating potent HIV-1 PR ligands often achieve large negative free energies primarily via polar and hydrophobic contacts [18]. Accordingly, a quinazoline inhibitor (CID-2135609) showed very stable binding and maintained key contacts with Asp25 and other residues during MD [18]. Similarly, BAN’s strong binding energy correlates with its robust hydrogen-bond network.

However, BAN’s binding conferred greater structural flexibility to the protease than 3TL. The backbone RMSD of the BAN-bound complex stabilized around 0.34 nm (Table 1, Fig. 2), substantially higher than 3TL’s ∼0.22 nm. Especially, Holo 3TL converged very quickly and maintained minimal fluctuation, indicating a highly stable conformation under MD. This behavior is akin to that reported for high-affinity inhibitors in which CID-2135609-bound PR showed minimal backbone deviation and conserved secondary structure [18]. By contrast, BAN induced larger RMSD oscillations, suggesting that its bound state allows the protein to deviate more from the starting structure. Residue RMSF analysis confirmed this. Accordingly, the BAN complex exhibits broad flexibility increases across loop and flap regions, residues 50–250, 750–950, and a peak near 1500. The average RMSF of BAN-bound PR was nearly double that of 3TL-bound. Such enhanced mobility is consistent with fewer stabilizing intraprotein constraints. In agreement, the Asp25 catalytic flaps likely sample wider conformational space when BAN is bound. In contrast, 3TL’s RMSF profile closely matched the apo enzyme, indicating 3TL rigidifies the enzyme much like CID-2135609 and other potent PIs do [16].

Despite BAN’s induced flexibility, all three systems had similar global SASA (63–66 nm²), implying no large-scale unfolding. The slight SASA increase in the BAN complex aligns with its elevated loop mobility. Protruding loops modestly expose more surface area without collapsing the structure. Particularly, previous work observed only minor SASA changes upon ligand binding for stable PR complexes [18]. Thus, BAN’s effect is subtle at the tertiary scale.

A key distinction between BAN and 3TL is their hydrogen-bonding patterns. Over 100 ns MD, 3TL formed only 13 transient H-bonds with PR (highest occupancies 0.7–0.8%), mainly to Gly48 and Phe53. In contrast, BAN established 18 H-bonds, including a highly persistent one with Asp25 (>100% occupancy, indicating multiple concurrent bonds) and stable interactions with Leu24 (20.7%), Gly48 (11.4%), and Gly49 (8.3%). This dense H-bond network likely underlies BAN’s favorable interaction energy. These findings are consonant with other studies. In this regard, residues Asp25, Gly27–30, and Ile50 often mediate PR inhibitor binding via H-bonds or van der Waals forces [37–39]. For example, CID-2135609 engaged Asp25 and Gly49 as key contacts [18]. The persistent Asp25 bonding by BAN is particularly encouraging, as this catalytic residue is a critical anchor in effective inhibitors. In contrast, 3TL’s lack of strong Asp25 bonding suggests it relies more on hydrophobic packing than on polar anchoring. Overall, BAN’s richer polar contacts hint at strong affinity despite the protein’s concomitant gain in flexibility.

Interestingly, BAN binding weakened several stabilizing intraprotein salt bridges. In the 3TL complex, the GLU34– LYS20 salt bridge (18.7% occupancy) and ASP30–LYS45 (10.2%) were the most persistent electrostatic contacts . These salt bridges likely help clamp PR’s folds. BAN reduced their occupancies drastically (GLU34–LYS20 to 3.9%; ASP30–LYS45 to 2.5%). The loss of these interactions is consistent with the higher RMSF and PCA variance observed for BAN-bound PR. To our knowledge, prior computational HIV-1 PR studies have seldom detailed salt-bridge dynamics; nonetheless, it is plausible that attenuating these electrostatic anchors contributes to the increased flap motion seen with BAN. Overall, BAN appears to induce a distinct conformational ensemble, trading intraprotein stability for stronger ligand contacts.

Principal component analysis provides further insight. Accordingly, the first four PCs of 3TL-bound PR captured 61.3% of variance (PC1 37.0%), whereas BAN-bound PR captured 72.1% (PC1 45.7%). The larger cumulative variance and trace, 3.49 vs 2.15, indicate more collective motion under BAN. The dominant mode corresponds to the flap “open-closed” motion. This suggests that BAN, despite binding tightly, permits more pronounced flap mobility than 3TL. The wider spread of trajectory points in the BAN PCs reinforces that BAN samples a broader conformational space. Such enhanced dynamics could affect inhibitor efficacy or residence time. Overall, 3TL binding confines the protease into a narrow, stable energy basin, while BAN induces a more adaptable state with deeper local minima, likely aided by its extensive H-bond network.

Taken together, this study indicates that BAN, HONH-BENZYLMALONYL-L-ALANYLGLYCINE-P-NITROANILIDE, is capable of binding HIV-1 PR with affinity comparable to 3TL, via both hydrophobic and strong polar interactions. However, BAN’s mechanism differs. It sustains the protease-ligand association by forming additional hydrogen bonds at Asp25 while simultaneously loosening intraprotein salt bridges and increasing overall flexibility. This contrasts with 3TL and other high-affinity PIs, which tend to “lock” PR into a rigid bound state. Thus, BAN represents a novel scaffold whose inhibition profile may be enthalpically driven. Future work should evaluate whether BAN’s dynamic binding leads to different inhibition kinetics or resistance profiles. The combined literature and simulation analyses underscore the importance of integrating multiple computational metrics to fully characterize protease–inhibitor complexes.

## Conclusions

Here, a DrugBank-derived compound, BAN, identified as a promising HIV-1 protease inhibitor through structure-based pharmacophore screening, docking, and MD simulation. Docking predicted BAN binds the protease with energy comparable to the reference inhibitor 3TL. Over 100 ns MD, BAN formed a robust hydrogen-bond network, including a highly persistent Asp25 interaction, and achieved a favorable total interaction energy. However, BAN also induced notably higher backbone and side-chain fluctuations in the enzyme compared to 3TL. BAN binding disrupted key intramolecular salt bridges and enlarged the conformational ensemble. In contrast, 3TL stabilized the protease in a rigid, low-mobility state with few lasting H-bonds. These findings suggest BAN’s binding is energetically favorable but dynamically distinct, balancing strong polar contacts with induced flexibility. The similar SASA across systems implies no unfolding, but the elevated mobility under BAN may influence inhibitor potency or resistance. Altogether, BAN emerges as a novel scaffold worthy of experimental validation. Its distinct binding signature, strong enthalpic interactions coupled with dynamic protein response, highlights a new chemotype for HIV-1 PR inhibition. Future optimization of BAN’s structure could yield high-affinity inhibitors that exploit its potent Asp25 interactions while enhancing structural rigidity. This study demonstrates the power of integrated computational design and analysis in uncovering and characterizing drug repurposing against HIV-1.

## Acknowledgments

Not applicable.

## Author contributions

A.M.R. Alrashedi designed the study, conducted the primary investigation, and wrote the original manuscript. A.S.K. Nazal performed the formal analysis and validated the data. L.A.R. Al-Iessa contributed to data visualization and provided essential resources. M.A.H.A. Rabeea supervised the project and assisted in reviewing and editing the manuscript. All authors reviewed the final manuscript and approved it for publication.

## Data availability statement

The datasets supporting the conclusions of this article are included within the article and its additional files.

## Ethics declarations

Not applicable.

## Consent to participate/Consent to publish

All authors have read and agreed to the published version of the manuscript.

## Generative AI statement

The author(s) declare that no Gen AI was used in the creation of this manuscript.

## References

1. Abraham, M. J., Murtola, T., Schulz, R., Páll, S., Smith, J. C., Hess, B., et al. (2015). Gromacs: High performance molecular simulations through multi-level parallelism from laptops to supercomputers. SoftwareX 1–2, 19–25. doi: 10.1016/j.softx.2015.06.001

2. Afgan, E., Nekrutenko, A., Grüning, B. A., Blankenberg, D., Goecks, J., Schatz, M. C., et al. (2022). The Galaxy platform for accessible, reproducible and collaborative biomedical analyses: 2022 update. Nucleic Acids Res 50, W345–W351. doi: 10.1093/nar/gkac247

3. Agniswamy, J., Kneller, D. W., Brothers, R., Wang, Y. F., Harrison, R. W., and Weber, I. T. (2019). Highly Drug-Resistant HIV-1 Protease Mutant PRS17 Shows Enhanced Binding to Substrate Analogues. ACS Omega 4, 8707–8719. doi: 10.1021/ACSOMEGA.9B00683/ASSET/IMAGES/LARGE/AO-2019-00683E_0005.JPEG

4. Arrigoni, R., Santacroce, L., Ballini, A., and Palese, L. L. (2023). AI-Aided Search for New HIV-1 Protease Ligands. Biomolecules 13. doi: 10.3390/BIOM13050858

5. Askari, F. S., Ebrahimi, M., Parhiz, J., Hassanpour, M., Mohebbi, A., and Mirshafiey, A. (2022). Digging for the discovery of SARS-CoV-2 nsp12 inhibitors: a pharmacophore-based and molecular dynamics simulation study. Future Virol. doi: 10.2217/fvl-2022-0054

6. Askari, F. S., and Mohebbi, A. (2025). The dynamic states of hepatitis B virus capsid monomers under the impact of different class of capsid-assembly modulators. Scientific Reports 2025 15:1 15, 1–21. doi: 10.1038/s41598-025-18339-6

7. Baassi, M., Moussaoui, M., Soufi, H., Rajkhowa, S., Sharma, A., Sinha, S., et al. (2023). Towards designing of a potential new HIV-1 protease inhibitor using QSAR study in combination with Molecular docking and Molecular dynamics simulations. PLoS One 18, e0284539. doi: 10.1371/JOURNAL.PONE.0284539

8. Baei, B., Askari, P., Askari, F. S., Kiani, S. J., and Mohebbi, A. (2025). Pharmacophore modeling and QSAR analysis of anti-HBV flavonols. PLoS One 20, e0316765. doi: 10.1371/journal.pone.0316765

9. Blankenberg, D., Coraor, N., Von Kuster, G., Taylor, J., and Nekrutenko, A. (2011). Integrating diverse databases into an unified analysis framework: A Galaxy approach. Database 2011. doi: 10.1093/database/bar011

10. Case, D. A., Aktulga, H. M., Belfon, K., Cerutti, D. S., Cisneros, G. A., Cruzeiro, V. W. D., et al. (2023). AmberTools. J Chem Inf Model 63, 6183–6191. doi: 10.1021/ACS.JCIM.3C01153/ASSET/IMAGES/LARGE/CI3C01153_0002.JPEG

11. Case, D. A., Cerutti, D. S., Cruzeiro, V. W. D., Darden, T. A., Duke, R. E., Ghazimirsaeed, M., et al. (2025). Recent Developments in Amber Biomolecular Simulations. J Chem Inf Model. doi: 10.1021/ACS.JCIM.5C01063/ASSET/IMAGES/LARGE/CI5C01063_0001.JPEG

12. Case, D. A., Cheatham, T. E., Darden, T., Gohlke, H., Luo, R., Merz, K. M., et al. (2005). The Amber biomolecular simulation programs. J Comput Chem 26, 1668–1688. doi: 10.1002/JCC.20290

13. Chuntakaruk, H., Hengphasatporn, K., Shigeta, Y., Aonbangkhen, C., Lee, V. S., Khotavivattana, T., et al. (2024). FMO-guided design of darunavir analogs as HIV-1 protease inhibitors. Scientific Reports 2024 14:1 14, 1–17. doi: 10.1038/s41598-024-53940-1

14. Dakshinamoorthy, A., Asmita, A., and Senapati, S. (2023). Comprehending the Structure, Dynamics, and Mechanism of Action of Drug-Resistant HIV Protease. ACS Omega 8, 9748–9763. doi: 10.1021/ACSOMEGA.2C08279/ASSET/IMAGES/LARGE/AO2C08279_0008.JPEG

15. Das, D., Koh, Y., Tojo, Y., Ghosh, A. K., and Mitsuya, H. (2009). Prediction of Potency of Protease Inhibitors Using Free Energy Simulations with Polarizable Quantum Mechanics-Based Ligand Charges and a Hybrid Water Model. J Chem Inf Model 49, 2851. doi: 10.1021/CI900320P

16. Deganutti, G., Prischi, F., and Reynolds, C. A. (2021). Supervised molecular dynamics for exploring the druggability of the SARS-CoV-2 spike protein. J Comput Aided Mol Des 35, 195–207. doi: 10.1007/s10822-020-00356-4

17. Eberhardt, J., Santos-Martins, D., Tillack, A. F., and Forli, S. (2021). AutoDock Vina 1.2.0: New Docking Methods, Expanded Force Field, and Python Bindings. J Chem Inf Model 61, 3891–3898. doi: 10.1021/ACS.JCIM.1C00203/SUPPL_FILE/CI1C00203_SI_002.ZIP

18. Falls, Z., Fine, J., Chopra, G., and Samudrala, R. (2022). Accurate Prediction of Inhibitor Binding to HIV-1 Protease Using CANDOCK. Front Chem 9, 775513. doi: 10.3389/FCHEM.2021.775513/BIBTEX

19. Martin, S. A., Cane, P. A., Pillay, D., and Mbisa, J. L. (2021). Coevolved Multidrug-Resistant HIV-1 Protease and Reverse Transcriptase Influences Integrase Drug Susceptibility and Replication Fitness. Pathogens 2021, *Vol.* 10*, Page* 1070 10, 1070. doi: 10.3390/PATHOGENS10091070

20. Mirarab, A., Mohebbi, A., Moradi, A., Javid, N., Vakili, M. A., and Tabarraei, A. (2017). Frequent pUL27 Variations in HIV-Infected Patients. Intervirology 59, 262–266. doi: 10.1159/000471484

21. Mohebbi, A. (2023). Ligand-based 3D pharmacophore modeling, virtual screening, and molecular dynamic simulation of potential smoothened inhibitors. J Mol Model 29. doi: 10.1007/s00894-023-05532-5

22. Mohebbi, A., Askari, F. S., Sammak, A. S., Ebrahimi, M., and Najafimemar, Z. (2021a). Druggability of cavity pockets within SARS-CoV-2 spike glycoprotein and pharmacophore-based drug discovery. Future Virol 16, 389–397. doi: 10.2217/fvl-2020-0394

23. Mohebbi, A., Askari, F. S., Sammak, A. S., Ebrahimi, M., and Najafimemar, Z. (2021b). Druggability of cavity pockets within SARS-CoV-2 spike glycoprotein and pharmacophore-based drug discovery. Future Virol 16, 389–397. doi: 10.2217/fvl-2020-0394

24. Mohebbi, A., Mirarab, A., Shaddel, R., Shafaei Fallah, M., and Memarian, A. (2021c). Molecular Dynamic Simulation and Docking of Cyclophilin A Mutants with its Potential Inhibitors. Journal of Clinical and Basic Research 5, 26–41. doi: 10.52547/jcbr.5.2.26

25. Mohebbi, A., Mohammadi, S., and Memarian, A. (2016). Prediction of HBF-0259 interactions with hepatitis B Virus receptors and surface antigen secretory factors. Virusdisease 27, 234–241. doi: 10.1007/s13337-016-0333-9

26. Mohebbi, A., Nabavi, S. P. T., Naderi, M., Sharifian, K., Behnezhad, F., Mohebbi, M., et al. (2025). Computer-aided drug repurposing & discovery for Hepatitis B capsid protein. In Silico Pharmacology 2025 13:1 13, 1–22. doi: 10.1007/S40203-025-00314-8

27. Nabavi, S. P. T., Askari, F. S., Askari, P., and Mohebbi, A. (2024). Molecular dynamic simulation of Cyclophilin A in complex with Sanglifehrin A. doi: 10.21203/RS.3.RS-5005208/V1

28. Namthabad, S., Appl, E. M.-I. J. R., and 2014, undefined (2014). Molecular Docking of HIV-1 Protease using Alkaloids from Tinospora cordifolia. International Journal of Research and Applications 1, 12–16. doi: 10.17812/ijra.1.1(3)2014

29. Okafor, S. N., Angsantikul, P., and Ahmed, H. (2022). Discovery of Novel HIV Protease Inhibitors Using Modern Computational Techniques. Int J Mol Sci 23. doi: 10.3390/IJMS232012149

30. Perryman, A. L., Zhang, Q., Soutter, H. H., Rosenfeld, R., McRee, D. E., Olson, A. J., et al. (2010). Fragment-Based Screen against HIV Protease. Chem Biol Drug Des 75, 257–268. doi: 10.1111/J.1747-0285.2009.00943.X

31. Rahman, M. M., Saha, T., Islam, K. J., Suman, R. H., Biswas, S., Rahat, E. U., et al. (2020). Virtual screening, molecular dynamics and structure–activity relationship studies to identify potent approved drugs for Covid-19 treatment. J Biomol Struct Dyn, 1–11. doi: 10.1080/07391102.2020.1794974

32. Rehan Ahmad, S., Zeyaullah, M., AlShahrani, A. M., Mohieldin, A., Alam, M. S., and Altijani, A. A. G. (2025). Structure-Based Identification and Molecular Characterization of CID-2135609 as a Potent Small Molecule Modulator of HIV-1 Protease. J Cell Biochem 126. doi: 10.1002/JCB.70058

33. Sadybekov, A. V., and Katritch, V. (2023). Computational approaches streamlining drug discovery. Nature 2023 616:7958 616, 673–685. doi: 10.1038/s41586-023-05905-z

34. Śledź, P., and Caflisch, A. (2018). Protein structure-based drug design: from docking to molecular dynamics. Curr Opin Struct Biol 48, 93–102. doi: 10.1016/j.sbi.2017.10.010

35. Sousa Da Silva, A. W., and Vranken, W. F. (2012). ACPYPE - AnteChamber PYthon Parser interfacE. BMC Res Notes 5, 1–8. doi: 10.1186/1756-0500-5-367/FIGURES/3

36. Trott, O., and Olson, A. J. (2009). AutoDock Vina: Improving the speed and accuracy of docking with a new scoring function, efficient optimization, and multithreading. J Comput Chem 31, NA-NA. doi: 10.1002/jcc.21334

37. Venkatachalam, S., Muralidharan, N., Pandian, R., Sayed, Y., and Gromiha, M. M. (2025). Integrative Computational Approaches for Understanding Drug Resistance in HIV-1 Protease Subtype C. Viruses 2025, *Vol.* 17*, Page* 850 17, 850. doi: 10.3390/V17060850

38. Volarath, P., Harrison, R., and Weber, I. (2007). Structure based drug design for HIV protease: from molecular modeling to cheminformatics. Curr Top Med Chem 7, 1030–1038. doi: 10.2174/156802607780906744

39. Weber, I. T., Kneller, D. W., and Wong-Sam, A. (2015). Highly resistant HIV-1 proteases and strategies for their inhibition. Future Med Chem 7, 1023. doi: 10.4155/FMC.15.44

